# Hot-wiring dynein-2 uncovers roles for IFT-A in retrograde train assembly and motility

**DOI:** 10.1101/2023.05.10.539247

**Authors:** Francisco Gonçalves-Santos, Ana R. G. De-Castro, Diogo R. M. Rodrigues, Maria J. G. De-Castro, Reto Gassmann, Carla M. C. Abreu, Tiago J. Dantas

## Abstract

Intraflagellar transport (IFT) trains, built around IFT-A and IFT-B complexes, are carried by opposing motors to import and export ciliary cargo. While transported by kinesin-2 on anterograde IFT trains, the dynein-2 motor adopts an autoinhibitory conformation until it needs to be activated at the ciliary tip to power retrograde IFT. Growing evidence has linked the IFT-A complex to retrograde IFT; however, its roles in this process remain unknown.

Here, we used CRISPR/Cas9-mediated editing to disable the dynein-2 autoinhibition mechanism in *Caenorhabditis elegans*, and assessed its impact on IFT with high-resolution live imaging and photobleaching analyses. Remarkably, this dynein-2 “hot-wiring” approach reignites retrograde motility inside IFT-A-deficient cilia, without triggering tug-of-war events. In addition to providing functional evidence that multiple mechanisms maintain dynein-2 inhibited during anterograde IFT, our data uncover key roles for IFT-A in: mediating motor-train coupling during IFT turnaround; promoting retrograde IFT initiation; and modulating dynein-2 retrograde motility.

**Highlights:**

- Hot-wiring mutations enable dynein-2 to undergo retrograde movement in IFT-A-deficient cilia
- Disabling dynein-2 autoinhibition reveals that multiple mechanisms restrain dynein-2 activity during anterograde IFT
- IFT-A promotes retrograde IFT initiation and efficient dynein-2 motility
- IFT-A mediates dynein-2 coupling to retrograde IFT trains

## INTRODUCTION

Cilia are microtubule-based projections specialized in various cellular functions, such as cellular locomotion, sensory perception, and signal transduction. The assembly and function of cilia depend on intraflagellar transport (IFT), a bidirectional transport system that uses large protein assemblies, known as IFT trains, to import and export ciliary cargo. The scaffold of IFT trains is formed by the multiprotein IFT-B and IFT-A complexes, which serve as binding platforms for adaptors and ciliary cargos (Nakayama and Katoh, 2020; Pigino, 2021).

IFT trains are transported along axonemal microtubules by two types of molecular motors: kinesin-2 and dynein-2. In some cilia, anterograde IFT is powered by a single class of kinesin-2 motors (heterotrimeric kinesin-2), while in others, such as in *Caenorhabditis elegans* neuronal cilia, the homodimeric kinesin-2 also comes into play, cooperating with the former to mediate train transport of ciliary cargos from the base to the tip (Prevo et al., 2017). Following IFT train remodeling at the ciliary tip, dynein-2 is activated to power the retrograde transport of cargos and retrieve the IFT machinery back to the ciliary base (Pigino, 2021). At the core of the dynein-2 complex are two force-producing heavy chains (HC, known as CHE-3/DHC-2 in *C. elegans* (Signor et al., 1999)). These bind accessory subunits such as intermediate and light intermediate chains that stabilize the motor and regulate its activity (IC and LIC, known in *C. elegans* as WDR-60 and XBX-1, respectively (De-Castro et al., 2022; Schafer et al., 2003)).

Given that kinesin-2 and dynein-2 are opposing motors, their activity must be tightly regulated to prevent resistance to movement (e.g. tug-of-war events) and maximize IFT efficiency (Mijalkovic et al., 2017). Indeed, recent studies have shown that the dynein-2 motor adopts an autoinhibitory/closed conformation, trapping the linker and microtubule-binding stalk regions within a motor-motor stacking interface (Figure 1A, 1B)(Jordan et al., 2018; Toropova et al., 2017; Toropova et al., 2019). Complementing this mechanism, the inactivated dynein-2 complex is loaded onto anterograde trains with its motor domains facing towards the ciliary membrane, which was proposed to further prevent it from engaging with the microtubule track while being transported by kinesin-2 (Jordan et al., 2018). Upon IFT turnaround, dynein-2 needs to be converted into an active state in order to initiate retrograde movement. However, the process through which this occurs remains unknown.

**Figure 1.**
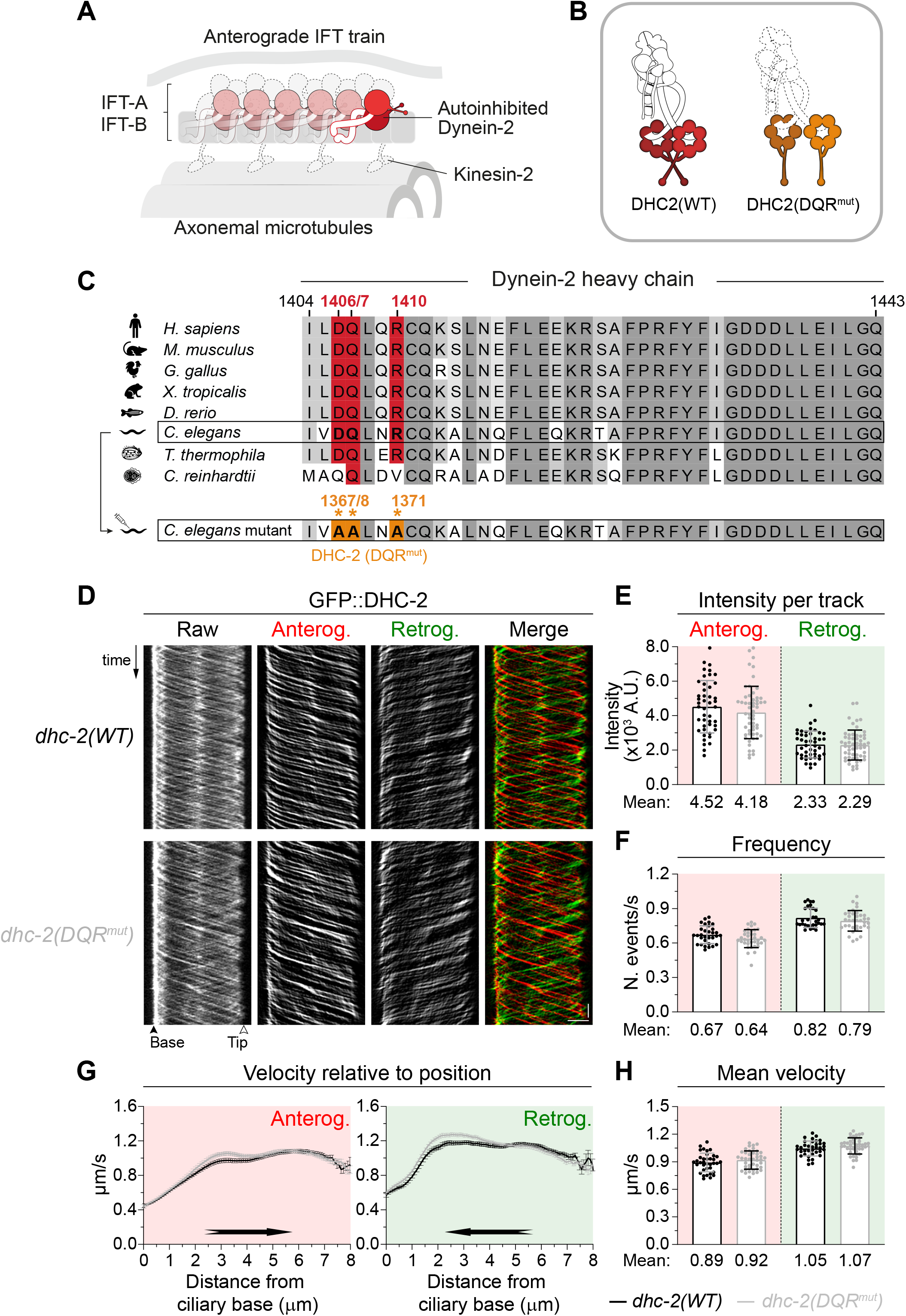
Mutating the conserved DQR residues of the dynein-2 autoinhibition interface does not alter IFT kinetics in *C. elegans* ciliay. **(A)** Illustration depicting the two concurrent mechanisms postulated to restrain dynein-2 activity during anterograde IFT: the autoinhibitory conformation of the motor (Toropova et al., 2017) and its orientation on anterograde trains (Jordan et al., 2018). IFT machinery drawn for context (not to scale). **(B)** Representation of the autoinhibited (trapped) structure of the human dynein-2 motor (DHC2(WT), red) (Toropova et al., 2019), and the untrapped motor conformation obtained in vitro upon mutation of the DQR residues (DHC2(DQR^mut^), orange) (Toropova et al., 2017). Dotted lines indicate lack of structural knowledge about the active dynein-2 complex. **(C)** Multiple sequence alignment of the dynein-2 HC shows evolutionary conservation of the DQR amino acids (red) in most species. Dynein-2 DQR to AAA mutations engineered in *C. elegans* are highlighted in orange. **(D)** Kymographs of endogenously-labeled wild-type and DQR mutant forms of dynein-2 HC. Software-tracked particles moving anterogradely and retrogradely are shown in red and green, respectively. **(E)** Relative amount of dynein-2 motors per IFT particle, inferred from the average GFP::DHC-2 intensity along anterograde and retrograde tracks in cilia of wild-type and dynein-2 DQR^mut^ animals (n≥2250 particles from ≥45 cilia). **(F)** IFT frequency measured by the number of IFT events per second detected at the middle segment of wild-type and dynein-2 DQR^mut^ cilia (n≥31 cilia). **(G)** Velocities of DHC-2 particles relative to their position along cilia of wild-type and dynein-2 DQR^mut^ animals. **(H)** Mean anterograde and retrograde IFT velocities in the indicated strains (n≥1800 particles from ≥36 cilia). XY graphs show mean ±SEM; column graphs show mean ±SD. Unpaired t-test was used to analyze datasets in E, F and H. Scale bar: vertical 5 s, horizontal 2 μm. See also Video S1.

Insights into retrograde IFT have emerged from research on the IFT-A complex, composed of six subunits: IFT144, IFT140, IFT122, IFT121, IFT139 and IFT43 (known in *C. elegans* as DYF-2/IFT-144, CHE-11/IFT-140, DAF-10/IFT-122, IFTA-1/IFT-121,

IFT-139, IFT-43, respectively (Blacque et al., 2006; Efimenko et al., 2006; Perkins et al., 1986; Scheidel and Blacque, 2018; Yi et al., 2017); Table S1; Figure S2A). Indeed, studies in diverse ciliated organisms have shown that loss of the IFT-A complex results in phenotypes reminiscent of dynein-2 mutants (Hesketh et al., 2022; Iomini et al., 2001; Nakayama and Katoh, 2020; Piperno et al., 1998; Quidwai et al., 2021; Taschner and Lorentzen, 2016). Yet, how IFT-A contributes to retrograde IFT has remained elusive.

Here, by genetically modifying the dynein-2 HC, we were able to “hot-wire” the motor and reignite its retrograde motility in IFT-A deficient cilia of *C. elegans* sensory neurons. Strikingly, the remaining IFT machinery was left stranded inside these cilia, providing clear evidence that the IFT-A complex mediates both retrograde train assembly and dynein-2 motility.

## RESULTS

### Mutating the conserved DQR residues of the dynein-2 autoinhibition interface does not alter IFT dynamics

To determine the importance of the dynein-2 autoinhibition mechanism in vivo, we sought to recreate in *C. elegans* the mutations that untrap the human dynein-2 HCs in vitro (Figure 1B). This consisted of mutating three highly conserved polar amino acids in the linker domain of the endogenous dynein-2 HC into three nonpolar amino acids (D1367A, Q1368A and R1371A in *C. elegans*; Figure 1C). In doing so, the resulting dynein-2 DQR^mut^ form becomes unable to establish the hydrogen bonds between the linker and the AAA4 module, shown to mediate the stacking interface of the autoinhibited dynein-2 (Toropova et al., 2017).

We then assessed the distribution and kinetics of the IFT machinery in cilia of phasmid neurons carrying the dynein-2 DQR mutations. Our analyses showed that dynein-2 DQR^mut^ cilia were of normal length (7.4±0.61µm versus 7.3±0.64µm in controls; n≥96 cilia), indicating that the DQR mutations do not affect cilia assembly, nor their elongation. Importantly, the distribution profiles of various IFT components in dynein-2 DQR^mut^ cilia almost perfectly matched those of the control strains (Figures S1A and S1B), suggesting that this mutant form of dynein-2 remains functional.

When examining IFT dynamics, we found that the frequency and average velocity of IFT in dynein-2 DQR^mut^ cilia remained unchanged in both directions (Figure 1D, 1F-1H, Video S1), without any detectable tug-of-war events between the two opposing IFT motors. Notably, the DQR mutations had only a minor impact on the amount of dynein-2 motors being loaded onto anterogradely moving particles (∼7.4% less), albeit not statistically significant, indicating that these mutations do not impair the packaging of the retrograde motor onto anterograde trains (Figure 1E). Additionally, dynein-2 DQR^mut^ animals successfully detected and avoided crossing a hyperosmotic glycerol ring as controls, reflecting a normal cilia-mediated response (Figure S1C).

Taken together, these results show that disrupting the autoinhibitory interface of dynein-2 does not compromise IFT or cilia function in *C. elegans*.

### DQR mutations hot-wire dynein-2, preventing its accumulation inside IFT-A-deficient cilia

Given that the DQR mutations promote the motility of human dynein-2 motor dimers in vitro (Toropova et al., 2017), we next sought to test whether these dynein-2 mutations would be able to compensate for retrograde IFT defects in vivo. We examined mutants of the IFT-A1/core module (Figure S2A; (Hesketh et al., 2022)), which result in prominent accumulations of the IFT machinery inside short and bulgy cilia in *C. elegans* (Albert et al., 1981; Efimenko et al., 2006; Perkins et al., 1986; Yi et al., 2017). We focused on the *ift-144(gk678), ift-140(tm3433)* and *ift-122(tm2878)* mutants, where we found that the integrity of the IFT-A complex was compromised, as evidenced by the almost complete loss of IFT-A2 subunits from their cilia (Figure S2).

Attesting to the importance of IFT-A for retrograde IFT, we found that the dynein-2 subunits DHC-2 and WDR-60 heavily accumulated inside cilia of *ift-144, ift-140* and *ift-122* mutants (Figure 2A-2D). Remarkably, combining these IFT-A1 mutants with the dynein-2 DQR mutations resulted in a clear rescue of DHC-2 and WDR-60 ciliary accumulations.

**Figure 2.**
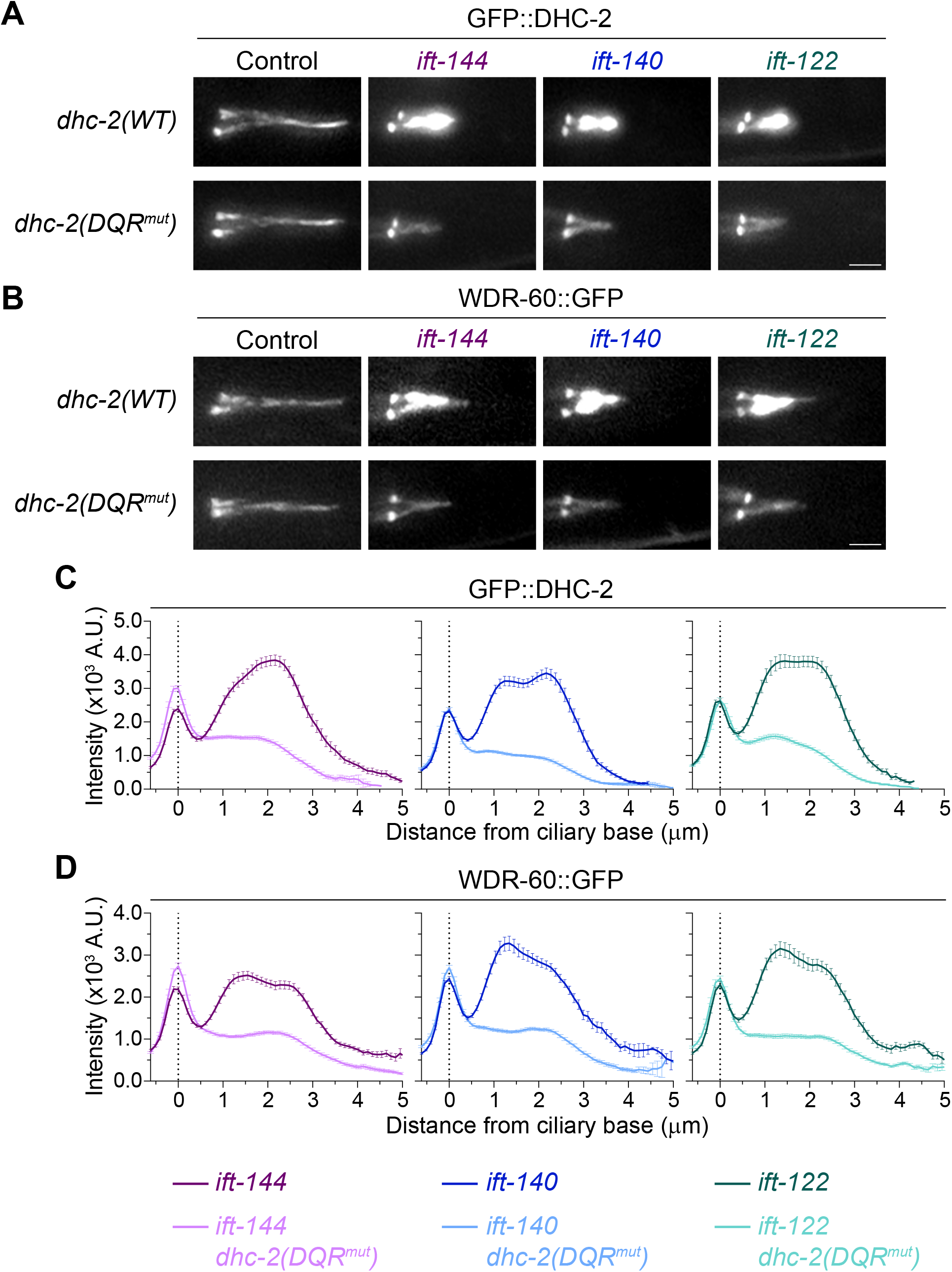
Promoting dynein-2 motor untrapping prevents its accumulation inside IFT-A1-deficient cilia. **(A)** Phasmid cilia from control and IFT-A1 mutant strains, expressing wild-type GFP::DHC-2 or GFP::DHC-2 DQR^mut^. **(B)** Phasmid cilia from control and IFT-A1 mutant strains, co-expressing WDR-60::GFP and the wild-type or DQR^mut^ form of DHC-2. **(C, D)** Distribution profile of GFP-tagged DHC-2 *(C)* and WDR-60 *(D)* along cilia of the indicated genotypes (n≥56 cilia). XY graphs show mean ±SEM. Scale bar: 2 μm.

These striking results suggest that the dynein-2 DQR mutations are capable of overcoming the requirement for IFT-A in retrograde dynein-2 motility. To test this hypothesis further, we performed live imaging and assessed the IFT kinetics of DHC-2(WT) and DHC-2(DQR^mut^) in IFT-A1 mutants (Figure 3A, 3B, Video S2). We found that DHC-2(WT) particles were still able to enter IFT-A1-deficient cilia and undergo anterograde IFT. DHC-2(WT) retrograde movement, on the other hand, was severely compromised, with hardly any retrograde tracks detected in these cilia. In fact, large amounts of static dynein-2 motors were observed in the kymographs of IFT-A-deficient cilia (Figure 3B). In sharp contrast, DHC-2(DQR^mut^) particles formed clear and continuous anterograde and retrograde tracks in all IFT-A1-deficient cilia, further advocating that the DQR mutations are able to reignite retrograde dynein-2 motility in the absence of IFT-A. Consistent with this rescue of retrograde initiation, virtually no DHC-2(DQR^mut^) particles were stalled inside IFT-A-deficient cilia (Figure 3B).

**Figure 3.**
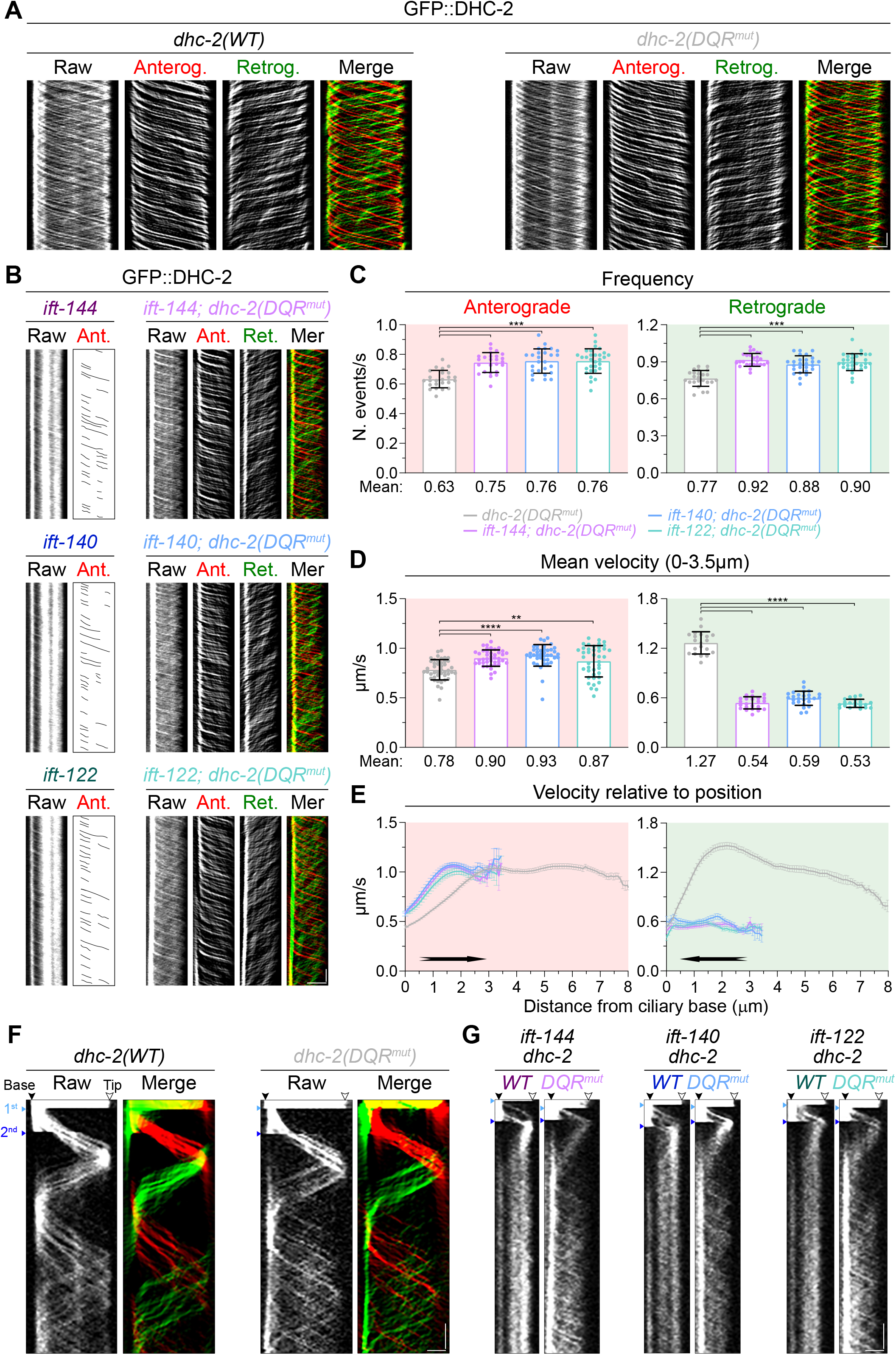
DQR mutations hot-wire dynein-2, restoring its motility in IFT-A1 mutant cilia. **(A, B)** Kymographs of the GFP-tagged wild-type and DQR^mut^ form of dynein-2, in control *(A)* and IFT-A1 mutant backgrounds *(B)*. Traces of particles moving anterogradely and retrogradely are shown in red and green, respectively. **(C)** Number of IFT events per second detected at the middle segment of IFT-A1-deficient cilia expressing GFP::DHC-2(DQR^mut^) (n≥24 cilia). **(D)** Mean IFT velocity of GFP::DHC-2(DQR^mut^) in control and IFT-A1 mutant backgrounds. **(E)** Velocity of GFP::DHC-2(DQR^mut^) particles at different positions along cilia in control and IFT-A1 mutant animals (n≥200 particle traces analyzed in ≥20 cilia). **(F, G)** Kymographs of the GFP-tagged wild-type and DQR^mut^ form of dynein-2, in control *(F)* and IFT-A1 mutant *(G)* backgrounds, upon sequential photobleaching (see also Figure S3B). The first and second photobleaching events indicated in the figure correspond to the bleaching of the signal inside the cilium body and the ciliary base, respectively. XY graphs show mean ±SEM; column graphs show mean ±SD. One-way ANOVA followed by the Holm–Sidak, Dunn, Tukey multiple comparisons tests were used to analyze the datasets in C and D, respectively. **, P≤0.01; ***, P≤0.001; ****, P≤0.0001. Scale bars: vertical 5 s, horizontal 2 μm. See also Videos S2-S4.

Through the analysis of the clearly distinguishable dynein-2 DQR^mut^ tracks, we found that the frequency of both anterograde and retrograde IFT events was increased in cilia from all IFT-A1 mutants (Figure 3C). Interestingly, while the velocity of anterograde IFT also significantly increased, we observed a striking reduction in retrograde IFT velocity (Figure 3D). Upon closer inspection, we found that anterogradely moving particles accelerated faster than controls, whereas retrograde movement was consistently slower throughout the shorter IFT-A1-deficient cilia, in spite of the presence of the dynein-2 DQR mutations (Figure 3E). Collectively, these observations reveal that, in addition to promoting retrograde IFT initiation, the IFT-A complex also contributes to the fast motility of dynein-2 during retrograde IFT. We also reason that the increased anterograde IFT kinetics associated with the loss of IFT-A might result from a reduction in the size of anterograde IFT trains, as they no longer carry IFT-A nor any components bound to them via IFT-A.

To further dissect the retrograde defects of IFT-A1 mutants, we performed sequential photobleaching experiments, which allows tracking of an isolated fraction of IFT particles as they travel along the axoneme of mutant cilia (Figure 3F, 3G, S3B). In agreement with the aforementioned experiments, the behavior of DHC-2(WT) and DHC-2(DQR^mut^) were very similar: particles reached the ciliary tip, turned around, and exited cilia in synchrony (Figure 3F, Video S3). In contrast, the majority of the DHC-2(WT) particles that entered IFT-A1-deficient cilia became stranded once they reached the ciliary tip, gradually being pushed away by the continuous influx of anterograde trains arriving at the tip. Remarkably, DHC-2(DQR^mut^) particles readily started retrograde movement after reaching the ciliary tip, successfully returning to the base of IFT-A1-deficient cilia (Figure 3G, Video S4). These results clearly show that the DQR mutations promote the initiation of retrograde dynein-2 motility in IFT-A1-deficient cilia, bypassing the requirement for IFT-A in this process.

### The IFT-A complex mediates dynein-2 coupling to retrograde IFT trains

Intriguingly, although the DQR mutations prevented dynein-2 accumulation inside IFT-A1-deficient cilia, ciliary elongation remained impaired (Figure S3A). This led us to hypothesize that, even in the presence of DHC-2(DQR^mut^), retrieval of the IFT machinery to the ciliary base might remain compromised in IFT-A1-deficient cilia. To investigate this, we analyzed the ciliary distribution of multiple IFT train components in IFT-A1 mutants, both with and without the dynein-2 DQR mutations. We started by examining the IFT-B complex, monitoring IFT-74 and IFT-54/DYF-11 of the IFT-B1 and IFT-B2 subcomplexes, respectively (Figure 4A, 4B). Both of these IFT-B components strongly accumulated inside cilia of the three IFT-A1 mutants, consistent with the defects in retrograde IFT observed before. Notably, expressing the DHC-2(DQR^mut^) form, which readily returns to the ciliary base, did not prevent the strong accumulation of IFT-B subunits at the tips of IFT-A1-deficient cilia (Figure 4A, 4B). These striking results reveal that IFT-A loss impairs the association of dynein-2 motors with IFT trains during retrograde IFT initiation.

**Figure 4.**
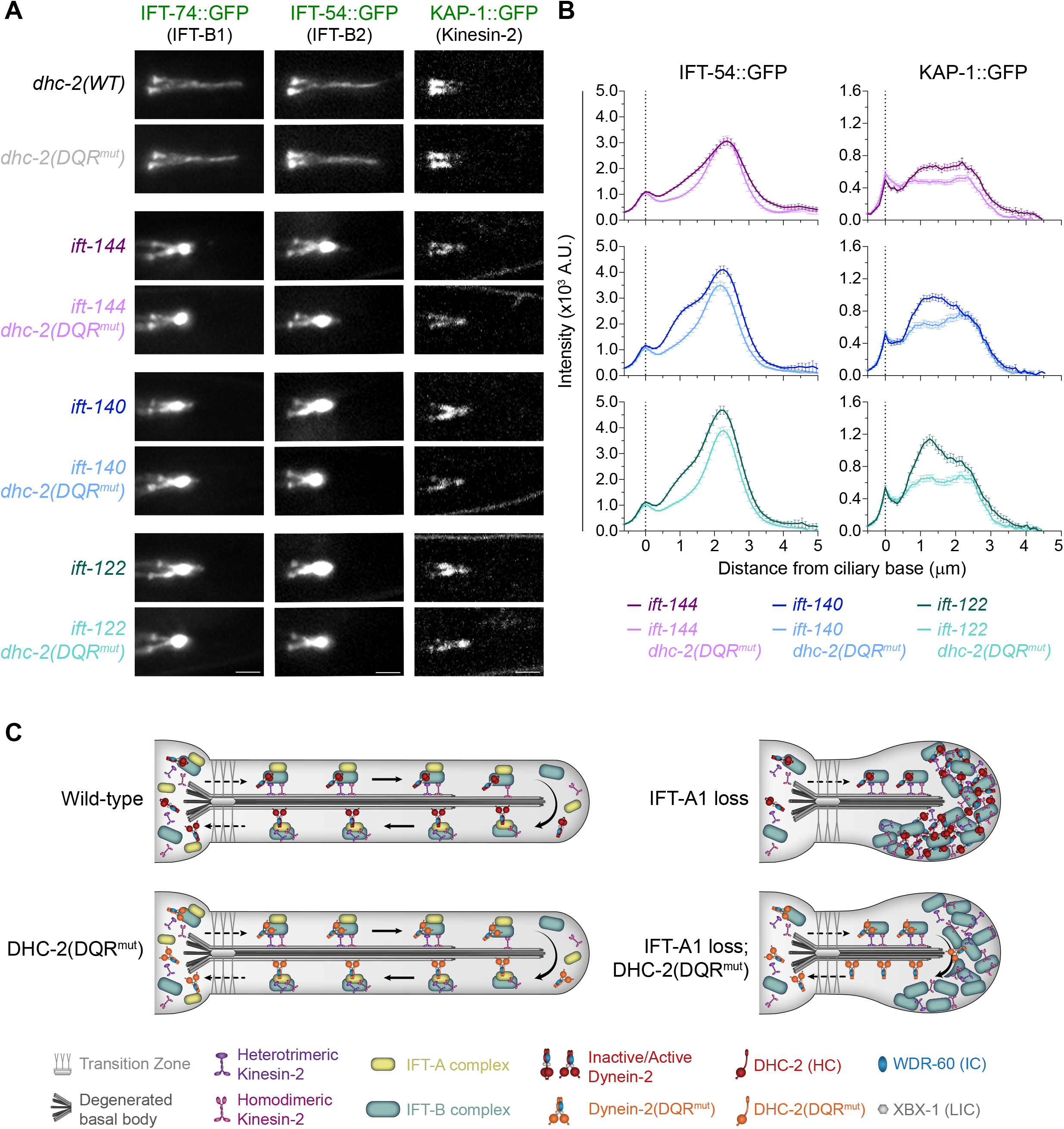
Loss of IFT-A disrupts the connection between dynein-2 and retrograde IFT trains. **(A)** Localization of representative components of the IFT machinery in phasmid cilia from control and IFT-A1 mutant strains, expressing the wild-type or DQR^mut^ form of DHC-2. Scale bars, 2 μm. **(B)** Distribution profile of IFT-54::GFP and KAP-1::GFP along cilia of the indicated genotypes (n≥62 cilia). XY graphs show mean ±SEM. **(C)** Model of IFT dynamics in cilia of the indicated genotypes. The IFT machinery of dynein-2 DQR^mut^ cilia behaves indistinguishably from wild-type. IFT-A1 loss strongly impairs retrograde IFT resulting in shorter cilia with severe accumulations of IFT components. The DHC-2 DQR mutations reignite dynein-2 motility at the ciliary tip of IFT-A1 mutants, restoring its ability to return to the ciliary base. The coupling of dynein-2 to retrograde IFT trains is compromised, leaving the remaining IFT machinery stranded inside IFT-A1-deficient cilia.

To further validate this, we analyzed the distribution of KAP-1 and OSM-3, subunits of the heterotrimeric and homodimeric kinesin-2 motors, respectively. We found that both anterograde motors also accumulated inside IFT-A1-deficient cilia, even in the presence of the dynein-2 DQR mutations, albeit at slightly reduced levels (Figure 4A, 4B, S4). These results further support our conclusion that IFT-A mediates the coupling of the retrograde motor to IFT trains upon their remodeling at the ciliary tip. Moreover, given that the BBSome is mostly absent from IFT-A-deficient cilia (Wei et al., 2012), our findings imply that the DQR mutations enable the dynein-2 motor complex to initiate retrograde movement on its own after IFT turnaround.

## DISCUSSION

In order to initiate retrograde IFT, dynein-2 motors need to overcome their regulatory constraints and transition into an active state capable of transporting IFT trains towards the ciliary base. In this study, we engineered a *C. elegans* dynein-2 DQR^mut^ strain, recapitulating the linker mutations that untrap the human dynein-2 motor and enhance its motility in vitro (Toropova et al., 2017). Using this powerful tool, we were able to dissect the poorly understood mechanisms regulating retrograde IFT and the specific contributions of IFT-A to this process.

### Complementary mechanisms restrain dynein-2 activity during anterograde IFT

Our analyses reveal that the *C. elegans* dynein-2 DQR^mut^ behaves essentially as wild-type, undergoing IFT in both directions with normal kinetics and similar rates of retrograde IFT initiation. Yet, we find that the same DQR mutations are able to overcome the requirement for IFT-A in the initiation of retrograde dynein-2 movement. This implies that the DQR mutations promote dynein-2 activation in vivo, most likely by untrapping its motor domains, as seen with recombinant dynein-2 DQR^mut^ motors in vitro (Toropova et al., 2017). Additionally, given that no tug-of-war events were detected in the dynein-2 DQR^mut^ and IFT turnaround occurred mostly at the ciliary tip, our data indicate that dynein-2 DQR^mut^ remains efficiently restrained during anterograde IFT.

Altogether, these results support that, besides the autoinhibition, additional mechanisms ensure that dynein-2 only becomes active upon IFT train remodelling and the need to initiate retrograde IFT. This conclusion is in line with recent findings in *Chlamydomonas reinhardtii* flagella, which showed that autoinhibited dynein-2 is carried on anterograde trains with its stalks oriented away from the microtubule track (Jordan et al., 2018). As proposed in the referred study, such positioning assures that dynein-2 is unable to engage with axonemal microtubules while being transported on anterograde trains.

Interestingly, our data also show that the DQR mutations that favor motor untrapping do not significantly interfere with dynein-2 loading onto anterograde trains. We speculate that the multiple links established between dynein-2 and anterograde IFT trains (Hiyamizu et al., 2023; Lacey et al., 2023; Pedersen et al., 2006; Toropova et al., 2019; Zhu et al., 2021) are capable of accommodating any conformational changes resulting from the DQR mutations.

Importantly, given that none of the IFT components examined accumulate inside dynein-2 DQR^mut^ cilia (Figure S1A, S1B), we conclude that hot-wiring dynein-2 alone does not interfere with the assembly of retrograde trains nor their transport towards the base.

### IFT-A1 subunits are essential for IFT-A complex assembly and its incorporation into cilia

Recent studies have revealed that the structure of the IFT-A complex is highly conserved across species (Hesketh et al., 2022; Jiang et al., 2023; Lacey et al., 2023; Ma et al., 2023; McCafferty et al., 2022; Meleppattu et al., 2022), aligning with its shared importance in retrograde IFT (Taschner and Lorentzen, 2016). In agreement with this, we show that the IFT-A1 mutants *ift-144(gk678), ift-140(tm3433)* and *ift-122(tm2878)* accumulate large amounts of IFT components inside their short and bulgy cilia. Consistent with the high severity of the phenotypes, we find that the integrity of the IFT-A complex is completely compromised in all of these IFT-A1 mutants, as judged by the impaired ciliary recruitment of other IFT-A subunits. We note, however, that a residual amount of IFT-139 was still detectable inside IFT-A1-deficient cilia (Figure S2B, S2C), consistent with previous observations in another *ift-140* mutant (Scheidel and Blacque, 2018). This result suggests that, even on its own, IFT-139 can establish weak connections with the anterograde IFT train that enable it to enter IFT-A-deficient cilia. This possibility is further supported by recent structural data on the IFT-A complex showing that IFT139 is in close contact with IFT-B subunits in anterograde IFT trains (Lacey et al., 2023; Ma et al., 2023; Meleppattu et al., 2022).

### IFT-A promotes retrograde IFT initiation and efficient dynein-2 motility

Our live imaging and sequential photobleaching data provide important insights into the functional roles of IFT-A in retrograde IFT. We show that, despite undergoing anterograde transport and reaching the ciliary tip, nearly all dynein-2 motors fail to move retrogradely in IFT-A mutants, becoming heavily accumulated inside their cilia. These data indicate that the IFT-A complex is instrumental for the robust initiation of dynein-2 retrograde movement. This conclusion is further supported by the fact that the hot-wiring DQR mutations are able to reignite retrograde dynein-2 movement in the absence of the IFT-A complex.

Notably, hot-wiring dynein-2 also allowed us to uncover another key contribution of IFT-A during retrograde movement. More specifically, our findings support a role for IFT-A in the regulation of dynein-2 motility, as evidenced by the consistently slower retrograde velocity of dynein-2 DQR^mut^ observed in all IFT-A mutants. As the DQR mutations alone do not affect motor dynamics, the slower dynein-2 velocity in IFT-A mutants can be attributed to the absence of IFT-A or the loss of other regulators imported by this IFT complex.

### IFT-A mediates dynein-2 coupling to retrograde IFT trains

Interestingly, we find that restoring retrograde dynein-2 motility in IFT-A mutants does not rescue cilia length. We attribute this to the fact that hot-wired dynein-2 is unable to retrieve retrograde IFT trains to the ciliary base in the absence of IFT-A, as evidenced by the persistent accumulation of IFT-B and kinesin-2 components inside cilia. This uncovers yet another critical role of the IFT-A complex in mediating the connection between dynein-2 and retrograde IFT trains during their remodelling at the ciliary tip. Although we note that IFT-B and kinesin-2 accumulations were slightly reduced inside dynein-2 DQR^mut^ cilia lacking IFT-A, we speculate that the continuous flow of dynein-2 through the overcrowded ciliary environment ends up pushing some of these components along as the motor exits cilia. Alternatively, it is conceivable that, during IFT train remodeling, some links can still form between dynein-2 and IFT-B in the absence of IFT-A. However, considering that IFT components still end up largely accumulated inside IFT-A-deficient cilia, such potential interactions would mostly fail to withstand the pulling forces exerted by hot-wired dynein-2.

Consistent with a role for IFT-A in mediating motor-train coupling during retrograde IFT, a previous study found that an *ift-144* point mutation led to the accumulation of IFT-B subunits and the OSM-3 motor at the ciliary tip, without blocking retrograde dynein-2 motility (Wei et al., 2012).

### Model for how IFT-A contributes to dynein-2-mediated retrograde IFT

Based on our data, we propose a model (Figure 4C) wherein IFT-A makes several key contributions to retrograde IFT: (A) mediating motor-train coupling during retrograde train assembly and transport; (B) promoting dynein-2 activation for retrograde IFT initiation; and (C) boosting dynein-2 motility for optimal retrograde velocity. These functions might be performed by direct binding of IFT-A to dynein-2, and/or the IFT-A-mediated import of regulatory factors into cilia, such as the BBSome, TULP adaptor proteins and kinases (Hesketh et al., 2022; Liem et al., 2012; Mukhopadhyay et al., 2010; Wei et al., 2012). In this regard, biochemical and in vivo assays have provided evidence of a connection between IFT-A and dynein-2 (Hiyamizu et al., 2023; Vuolo et al., 2018; Wei et al., 2012; Yi et al., 2017). However, it is not clear if such interactions are direct, nor is it possible to distinguish whether these occur in the context of anterograde or retrograde IFT trains.

Given that the DQR untrapping mutations enable the dynein-2 motor to initiate retrograde motility in the absence of IFT-A, we speculate that IFT-A promotes dynein-2 activation by favoring the open/untrapped motor conformation. This leads us to propose that IFT-A promotes retrograde IFT initiation in a conceptually analogous way to dynactin and/or Lis1, which stabilize the open conformation of the cytoplasmic dynein-1 motor for activation of motility and cargo-binding (Zhang et al., 2017).

## ACKNOWLEDGEMENTS

We thank Dr Ana Carvalho for sharing equipment and for exchanging ideas. We are also very grateful to Dr Erwin Peterman and Dr Guangshuo Ou for providing *C. elegans* strains, and Dr Anthony Roberts for discussions.

This work was supported by the projects 2022.01955.PTDC (to T.J.D.) and 2022.01964.PTDC (to C.M.C.A.) from Fundação para a Ciência e a Tecnologia (FCT).

A.R.G.D. and D.R.M.R received PhD fellowships from FCT (UI/BD/152865/2022 and SFRH/BD/143985/2019, respectively) and support from the Molecular and Cell Biology, and the Biomedical Sciences PhD programs at ICBAS.

R.G, C.M.C.A., and T.J.D. salaries were also supported by FCT: CEECIND/00333/2017, CEECIND/01985/2018 and CEECIND/00771/2017, respectively.

The authors also thank the National Bioresource Project for *C. elegans* and the *Caenorhabditis* Genetics Center (CGC) for providing strains.

The authors declare no competing financial interests.

## AUTHOR CONTRIBUTIONS

F.G. and A.R.G.D. performed most of the experiments, actively contributed to the experimental design, analyzed data, helped to prepare figures and to write the manuscript. and D.R.M.R. helped with several experiments, quantifications and figure preparation. M.J.G.D. helped with some experiments and figure preparation. R.G. provided strains, reagents, equipment, and helped with experimental design. T.J.D. and C.M.C.A. conceived and supervised the project, designed and helped with experiments, analyzed and interpreted data, and helped to prepare figures and to write the manuscript. All authors read the manuscript and provided input for the final version.

## SUPPLEMENTAL INFORMATION

**Figures S1-S4** and **Table S1:** included in separate file.

**Video S1**. Live imaging of endogenously-labeled GFP::DHC-2, wild-type (top) and DQR mutant (bottom), undergoing IFT in *C. elegans* phasmid cilia. Images were acquired at 3 fps and playback is set at 15 fps (5x speed). Scale bar: 2 μm. Time is displayed as min:s.

**Video S2**. Live imaging of endogenously-labeled GFP::DHC-2, wild-type (top) and DQR mutant (bottom), in the three independent IFT-A1 mutant backgrounds. Images were acquired at 3 fps and playback is set at 15 fps (5x speed). Scale bar: 2 μm. Time is displayed as min:s.

**Video S3**. Sequential photobleaching of endogenously-labeled GFP::DHC-2, wild-type (top) and DQR mutant (bottom). By first photobleaching the cilium body, and then the ciliary base, the behavior of isolated IFT particles could be tracked as they traveled along cilia. Images were acquired at 3 fps and playback is set at 15 fps (5x speed). Scale bar: 2 μm. Time is displayed as min:s.

**Video S4**. Sequential photobleaching of endogenously-labeled GFP::DHC-2, wild-type (top) and DQR mutant (bottom), in the three independent IFT-A1 mutant backgrounds. By first photobleaching the cilium body, and then the ciliary base, the behavior of isolated IFT particles could be tracked as they traveled along cilia. Images were acquired at 3 fps and playback is set at 15 fps (5x speed). Scale bar: 2 μm. Time is displayed as min:s.

## METHODS

### RESOURCE AVAILABILITY

#### Lead contact and materials availability

Further information and requests for resources and reagents should be directed to and will be fulfilled by the Lead Contact, Tiago J. Dantas.

#### Data and code availability

All data generated and analyzed during this study are available from the Lead Contact upon reasonable request.

### EXPERIMENTAL MODEL AND SUBJECT DETAILS

#### Maintenance and generation of *Caenorhabditis elegans* strains

*C. elegans* strains were maintained at 20°C on standard nematode growth medium (NGM) plates seeded with bacteria (*Escherichia coli* OP50). Hermaphrodite *C. elegans* were used to maintain stable homozygous backgrounds as they have the ability to self-fertilize. To combine various genetic backgrounds, strains were crossed using standard procedures (Brenner, 1974) and mutant alleles were tracked by PCR-based genotyping. Some alternative *C. elegans* gene nomenclature is used throughout this study to facilitate its interpretation by readers working with other ciliated organisms, thereby leading to more transferable knowledge in the cilia field. Importantly, the alternative gene names used were approved by WormBase curators prior to the submission of this manuscript (changes will be made public in WormBase release WS289). See Table S1 for details.

## METHOD DETAILS

### Protein sequence alignments

Dynein-2 heavy chain sequences used in the alignments shown in Fig. 1C were obtained from the UniProt database, and have the following accession numbers: *Homo sapiens* - Q8NCM8; *Mus musculus* - Q45VK7; *Gallus gallus* - A0A8V0XK27; *Xenopus tropicalis* - A0A6I8PXK3; *Danio rerio* - A0A8M9PCL1; *Caenorhabditis elegans* - Q19542; *Tetrahymena thermophila* - Q5U9X1; *Chlamydomonas reinhardtii* - Q9SMH5. Jalview software (version 2.11.2.6) was used to align the protein sequences (Waterhouse et al., 2009), with the Muscle algorithm (Edgar, 2004).

### CRISPR/Cas9-mediated genome editing

Genetic loci of several genes encoding IFT components were edited by CRISPR/Cas9 using microinjection. Briefly, mixtures of Cas9 protein, guide RNAs, and homology templates consisting of DNA oligos or partially single-stranded DNA fragments were microinjected into the gonads of young adult hermaphrodites as described (Dokshin et al., 2018). Co-injection of a dominant marker aided in the selection of progeny. The presence of the desired alleles in the offspring was confirmed by PCR genotyping. The newly edited strains were outcrossed 4 to 5 times to ensure the absence of potential CRISPR/Cas9 off-target mutations.

### Fluorescence imaging

All imaging was carried out in phasmid cilia from young adult hermaphrodites expressing fluorescently-labeled IFT components, in temperature-controlled rooms kept at 20°C. Prior to imaging, animals were immobilized with 10 mM Levamisol, and placed on a 5% agarose pad mounted on a glass slide.

Imaging of fluorescently-labeled IFT components expressed at endogenous levels was carried out using an epifluorescence Axio Observer microscope (Zeiss), equipped with a Plan-Apochromat 63x/1.46 NA oil objective lens, an Orca Flash 4.0 camera (Hamamatsu), and controlled by ZEN software (Zeiss). Alternatively, cilia were imaged on an Olympus IX81 (Olympus, UK) inverted microscope equipped with an UPLSAPO 100x/1.40 NA oil objective lens, and an Andor Revolution XD spinning disk confocal system composed of a solid-state laser combiner (ALC-UVP 350i, Andor Technology, UK), a CSU-X1 confocal scanner (Yokogawa Electric Corporation), and an iXon^EM^+ DU-897 with 2x port coupler camera (Andor Technology), controlled by Andor IQ3 software (Andor Technology). Z-stacks were acquired with 0.4 µm spacing between each z-section.

Time-lapse imaging of IFT was performed using the same Andor Revolution XD spinning disk confocal system described above. Only phasmid cilia fully in focus in a single z-plane were selected for live imaging. 200 frames of each phasmid cilium were recorded at a rate of 3 frames/sec (333 ms per frame).

Sequential photobleaching of GFP::DHC-2 was performed during time-lapse imaging on a different Andor Revolution XD spinning disk confocal system (Andor Technology) equipped with a FRAPPA photobleaching module (Andor Technology). This system was mounted on an inverted microscope (Ti-E, Nikon), with solid-state lasers of 405 nm, 488 nm and 561 nm, a confocal scanner unit (CSU-X1; Yokogawa Electric Corporation) and an iXon^EM^+ camera (Andor Technology), all controlled by the Andor IQ3 software (Andor Technology). Using the 405 nm laser (60 mW) with 70% power and a 60x/1.40 NA oil-immersion Plan-Apochromat objective, an initial sweep was performed to the area corresponding to the cilium body in order to photobleach all fluorescence signal inside the axoneme. After allowing a few IFT particles to enter the cilium (approximately 3 seconds), the area of its base was also photobleached. In doing so, we were able to better follow the behavior of an individualized set of particles over time, even in cilia with strong accumulations of IFT components. Time-lapse image acquisition was also performed at a rate of 3 frames/sec (333 ms per frame).

### Image processing and analyses of live IFT

Z-stack and time-lapse acquisitions were processed and analyzed with Fiji software (Image J version 2.1.0/1.52v). The distribution profiles of IFT components were determined by drawing a line along the length of each cilium, from the base to the tip, and measuring the fluorescence intensity of the pixels along that line. Background signal was measured in a region next to the cilium and subtracted from the raw intensities obtained in the prior step. The resulting intensities determined along different cilia were averaged and plotted relative to the distance from the ciliary base.

The signal profile of GFP::DHC-2 was used to measure the length of cilia in wild-type and mutant strains.

Anterograde and retrograde IFT kymographs were generated in ImageJ (NIH) using the KymographClear toolset plugin (version 2.0; (Mangeol et al., 2016)). To ensure the quality of the data used to analyze IFT dynamics, only cilia completely visible in a single focal plane were used to generate the kymographs. The KymographDirect software (version 2.1; (Mangeol et al., 2016)) was then used to analyze IFT dynamics, automatically taking into account background and bleaching. The tracks of individual IFT particles moving along each cilium in either direction were automatically detected by the program as shown by (Mijalkovic et al., 2017), and further validated manual inspection. Reduced cilia size hampered the accuracy of the KymographDirect software, particularly when detecting retrograde tracks. Thus, the retrograde tracks of IFT-A mutant cilia expressing the dynein-2 DQR mutations were traced manually. Continuous IFT tracks drawn in the prior steps were then used to extract IFT velocities at different positions along cilia using the same software. Given the shorter size of IFT-A deficient cilia, the average velocities shown in Figure 3 were determined for the initial 3.5 µm of all cilia, thus facilitating a more direct comparison between mutants and control.

The frequency of anterograde and retrograde IFT events (data in Figures 1, 3) was determined using the method described by (Mijalkovic et al., 2017). More specifically, a vertical straight line was drawn across the same position in the ciliary middle segment of each anterograde or retrograde kymograph. The line drawn on each kymograph was then used to generate an intensity profile plot, where the number of intensity spikes were scored to determine the number of IFT events in each direction. This quantification reflects the number of distinguishable GFP::DHC-2 particles that move in each direction over time.

The average signal intensities of anterogradely or retrogradely moving GFP::DHC-2 particles were determined using KymographDirect as described (Mijalkovic et al., 2017). The software automatically measured the intensity of all pixels composing the track of each IFT particle. Each point of GFP::DHC-2 intensity plotted in Figure 1 corresponds to the average of all tracks (a minimum of 50), for either anterograde or retrograde IFT, from a particular cilium. At least 45 cilia were used per strain to accurately determine the average GFP::DHC-2 intensity particles moving in either direction.

### Osmotic Avoidance Assay

Osmotic avoidance assays were performed on NGM non-seeded plates at room temperature (∼20ºC), following the guidelines of (Sanders et al., 2015). Briefly, five young adult hermaphrodites were placed inside a glycerol ring with a diameter of approximately 1 cm, freshly prepared with a tube dipped in a 59% glycerol solution. Worm behavior was recorded for 5 min with a camera (QIClick digital CCD, QImaging) coupled to a stereomicroscope to determine whether they avoided crossing the glycerol ring. Worms that left the ring or stayed in contact with its glycerol border for more than 20 s were classified as escapers. Each repeat was carried out using worms of each strain carefully isolated before the experiment. Wild-type and *ift-54/dyf-11 (pe554)* worms were used as controls.

## QUANTIFICATION AND STATISTICAL ANALYSIS

### Data analyses and statistics

Statistical analyses of datasets were performed using GraphPad Prism software (version 9). Normality tests (Anderson-Darling test; D’Agostino-Pearson test; Shapiro-Wilk test; Kolmogorov-Smirnov test) were initially performed for each dataset, in order to determine if they followed a Gaussian distribution, thereby dictating the use of parametric or nonparametric statistical analyses. Pairwise comparisons were made using the two-tailed Student’s *t* test and the Mann-Whitney *U* tests for parametric and nonparametric data, respectively. Multiple comparisons were made using the parametric one-way ANOVA or the non-parametric Kruskal-Wallis test, followed by multiple comparisons tests. Differences were considered statistically significant for P values equal and below 5% (*P≤0.05; **P≤0.01; ***P≤0.001; ****P≤0.0001). XY position-averaged velocity and intensity distribution graphs are shown as mean ±SEM. Column graphs are shown as mean ± SD.

## REFERENCES

Albert, P.S., Brown, S.J., and Riddle, D.L. (1981). Sensory control of dauer larva formation in Caenorhabditis elegans. J Comp Neurol 198, 435–451. 10.1002/cne.901980305.

Blacque, O.E., Li, C., Inglis, P.N., Esmail, M.A., Ou, G., Mah, A.K., Baillie, D.L., Scholey, J.M., and Leroux, M.R. (2006). The WD repeat-containing protein IFTA-1 is required for retrograde intraflagellar transport. Mol Biol Cell 17, 5053–5062. 10.1091/mbc.e06-06-0571.

Brenner, S. (1974). The genetics of Caenorhabditis elegans. Genetics 77, 71–94.

De-Castro, A.R.G., Rodrigues, D.R.M., De-Castro, M.J.G., Vieira, N., Vieira, C., Carvalho, A.X., Gassmann, R., Abreu, C.M.C., and Dantas, T.J. (2022). WDR60-mediated dynein-2 loading into cilia powers retrograde IFT and transition zone crossing. J Cell Biol 221. 10.1083/jcb.202010178.

Dokshin, G.A., Ghanta, K.S., Piscopo, K.M., and Mello, C.C. (2018). Robust Genome Editing with Short Single-Stranded and Long, Partially Single-Stranded DNA Donors in Caenorhabditis elegans. Genetics 210, 781–787. 10.1534/genetics.118.301532.

Edgar, R.C. (2004). MUSCLE: multiple sequence alignment with high accuracy and high throughput. Nucleic Acids Res 32, 1792–1797. 10.1093/nar/gkh340.

Efimenko, E., Blacque, O.E., Ou, G., Haycraft, C.J., Yoder, B.K., Scholey, J.M., Leroux, M.R., and Swoboda, P. (2006). Caenorhabditis elegans DYF-2, an orthologue of human WDR19, is a component of the intraflagellar transport machinery in sensory cilia. Mol Biol Cell 17, 4801–4811. 10.1091/mbc.e06-04-0260.

Hesketh, S.J., Mukhopadhyay, A.G., Nakamura, D., Toropova, K., and Roberts, A.J. (2022). IFT-A structure reveals carriages for membrane protein transport into cilia. Cell 185, 4971–4985 e4916. 10.1016/j.cell.2022.11.010.

Hiyamizu, S., Qiu, H., Vuolo, L., Stevenson, N.L., Shak, C., Heesom, K.J., Hamada, Y., Tsurumi, Y., Chiba, S., Katoh, Y., et al. (2023). Multiple interactions of the dynein-2 complex with the IFT-B complex are required for effective intraflagellar transport. J Cell Sci 136. 10.1242/jcs.260462.

Iomini, C., Babaev-Khaimov, V., Sassaroli, M., and Piperno, G. (2001). Protein particles in Chlamydomonas flagella undergo a transport cycle consisting of four phases. J Cell Biol 153, 13–24. 10.1083/jcb.153.1.13.

Jiang, M., Palicharla, V.R., Miller, D., Hwang, S.H., Zhu, H., Hixson, P., Mukhopadhyay, S., and Sun, J. (2023). Human IFT-A complex structures provide molecular insights into ciliary transport. Cell Res 33, 288–298. 10.1038/s41422-023-00778-3.

Jordan, M.A., Diener, D.R., Stepanek, L., and Pigino, G. (2018). The cryo-EM structure of intraflagellar transport trains reveals how dynein is inactivated to ensure unidirectional anterograde movement in cilia. Nat Cell Biol 20, 1250–1255. 10.1038/s41556-018-0213-1.

Lacey, S.E., Foster, H.E., and Pigino, G. (2023). The molecular structure of IFT-A and IFT-B in anterograde intraflagellar transport trains. Nat Struct Mol Biol. 10.1038/s41594-022-00905-5.

Liem, K.F., Jr., Ashe, A., He, M., Satir, P., Moran, J., Beier, D., Wicking, C., and Anderson, K.V. (2012). The IFT-A complex regulates Shh signaling through cilia structure and membrane protein trafficking. J Cell Biol 197, 789–800. 10.1083/jcb.201110049.

Ma, Y., He, J., Li, S., Yao, D., Huang, C., Wu, J., and Lei, M. (2023). Structural insight into the intraflagellar transport complex IFT-A and its assembly in the anterograde IFT train. Nat Commun 14, 1506. 10.1038/s41467-023-37208-2.

Mangeol, P., Prevo, B., and Peterman, E.J. (2016). KymographClear and KymographDirect: two tools for the automated quantitative analysis of molecular and cellular dynamics using kymographs. Molecular biology of the cell 27, 1948–1957. 10.1091/mbc.E15-06-0404.

McCafferty, C.L., Papoulas, O., Jordan, M.A., Hoogerbrugge, G., Nichols, C., Pigino, G., Taylor, D.W., Wallingford, J.B., and Marcotte, E.M. (2022). Integrative modeling reveals the molecular architecture of the intraflagellar transport A (IFT-A) complex. Elife 11. 10.7554/eLife.81977.

Meleppattu, S., Zhou, H., Dai, J., Gui, M., and Brown, A. (2022). Mechanism of IFT-A polymerization into trains for ciliary transport. Cell 185, 4986–4998 e4912. 10.1016/j.cell.2022.11.033.

Mijalkovic, J., Prevo, B., Oswald, F., Mangeol, P., and Peterman, E.J. (2017). Ensemble and single-molecule dynamics of IFT dynein in Caenorhabditis elegans cilia. Nat Commun 8, 14591. 10.1038/ncomms14591.

Mukhopadhyay, S., Wen, X., Chih, B., Nelson, C.D., Lane, W.S., Scales, S.J., and Jackson, P.K. (2010). TULP3 bridges the IFT-A complex and membrane phosphoinositides to promote trafficking of G protein-coupled receptors into primary cilia. Genes Dev 24, 2180–2193. 10.1101/gad.1966210.

Nakayama, K., and Katoh, Y. (2020). Architecture of the IFT ciliary trafficking machinery and interplay between its components. Crit Rev Biochem Mol Biol 55, 179–196. 10.1080/10409238.2020.1768206.

Pedersen, L.B., Geimer, S., and Rosenbaum, J.L. (2006). Dissecting the molecular mechanisms of intraflagellar transport in chlamydomonas. Curr Biol 16, 450–459. 10.1016/j.cub.2006.02.020.

Perkins, L.A., Hedgecock, E.M., Thomson, J.N., and Culotti, J.G. (1986). Mutant sensory cilia in the nematode Caenorhabditis elegans. Dev Biol 117, 456–487. 10.1016/0012-1606(86)90314-3.

Pigino, G. (2021). Intraflagellar transport. Curr Biol 31, R530–R536. 10.1016/j.cub.2021.03.081.

Piperno, G., Siuda, E., Henderson, S., Segil, M., Vaananen, H., and Sassaroli, M. (1998). Distinct mutants of retrograde intraflagellar transport (IFT) share similar morphological and molecular defects. J Cell Biol 143, 1591–1601. 10.1083/jcb.143.6.1591.

Prevo, B., Scholey, J.M., and Peterman, E.J.G. (2017). Intraflagellar transport: mechanisms of motor action, cooperation, and cargo delivery. FEBS J 284, 2905–2931. 10.1111/febs.14068.

Quidwai, T., Wang, J., Hall, E.A., Petriman, N.A., Leng, W., Kiesel, P., Wells, J.N., Murphy, L.C., Keighren, M.A., Marsh, J.A., et al. (2021). A WDR35-dependent coat protein complex transports ciliary membrane cargo vesicles to cilia. Elife 10. 10.7554/eLife.69786.

Sanders, A.A., Kennedy, J., and Blacque, O.E. (2015). Image analysis of Caenorhabditis elegans ciliary transition zone structure, ultrastructure, molecular composition, and function. Methods Cell Biol 127, 323–347. 10.1016/bs.mcb.2015.01.010.

Schafer, J.C., Haycraft, C.J., Thomas, J.H., Yoder, B.K., and Swoboda, P. (2003). XBX-1 encodes a dynein light intermediate chain required for retrograde intraflagellar transport and cilia assembly in Caenorhabditis elegans. Mol Biol Cell 14, 2057–2070. 10.1091/mbc.e02-10-0677.

Scheidel, N., and Blacque, O.E. (2018). Intraflagellar Transport Complex A Genes Differentially Regulate Cilium Formation and Transition Zone Gating. Curr Biol 28, 3279–3287 e3272. 10.1016/j.cub.2018.08.017.

Signor, D., Wedaman, K.P., Orozco, J.T., Dwyer, N.D., Bargmann, C.I., Rose, L.S., and Scholey, J.M. (1999). Role of a class DHC1b dynein in retrograde transport of IFT motors and IFT raft particles along cilia, but not dendrites, in chemosensory neurons of living Caenorhabditis elegans. J Cell Biol 147, 519–530. 10.1083/jcb.147.3.519.

Taschner, M., and Lorentzen, E. (2016). The Intraflagellar Transport Machinery. Cold Spring Harb Perspect Biol 8. 10.1101/cshperspect.a028092.

Toropova, K., Mladenov, M., and Roberts, A.J. (2017). Intraflagellar transport dynein is autoinhibited by trapping of its mechanical and track-binding elements. Nat Struct Mol Biol 24, 461–468. 10.1038/nsmb.3391.

Toropova, K., Zalyte, R., Mukhopadhyay, A.G., Mladenov, M., Carter, A.P., and Roberts, A.J. (2019). Structure of the dynein-2 complex and its assembly with intraflagellar transport trains. Nat Struct Mol Biol 26, 823–829. 10.1038/s41594-019-0286-y.

Vuolo, L., Stevenson, N.L., Heesom, K.J., and Stephens, D.J. (2018). Dynein-2 intermediate chains play crucial but distinct roles in primary cilia formation and function. Elife 7. 10.7554/eLife.39655.

Waterhouse, A.M., Procter, J.B., Martin, D.M., Clamp, M., and Barton, G.J. (2009). Jalview Version 2--a multiple sequence alignment editor and analysis workbench. Bioinformatics 25, 1189–1191. 10.1093/bioinformatics/btp033.

Wei, Q., Zhang, Y., Li, Y., Zhang, Q., Ling, K., and Hu, J. (2012). The BBSome controls IFT assembly and turnaround in cilia. Nat Cell Biol 14, 950–957. 10.1038/ncb2560.

Yi, P., Li, W.J., Dong, M.Q., and Ou, G. (2017). Dynein-Driven Retrograde Intraflagellar Transport Is Triphasic in C. elegans Sensory Cilia. Curr Biol 27, 1448–1461 e1447. 10.1016/j.cub.2017.04.015.

Zhang, K., Foster, H.E., Rondelet, A., Lacey, S.E., Bahi-Buisson, N., Bird, A.W., and Carter, A.P. (2017). Cryo-EM Reveals How Human Cytoplasmic Dynein Is Auto-inhibited and Activated. Cell 169, 1303–1314 e1318. 10.1016/j.cell.2017.05.025.

Zhu, X., Wang, J., Li, S., Lechtreck, K., and Pan, J. (2021). IFT54 directly interacts with kinesin-II and IFT dynein to regulate anterograde intraflagellar transport. EMBO J 40, e105781. 10.15252/embj.2020105781.

