## Supplemental Figures S1-S4 and Table S1 for "Hot-wiring dynein-2 uncovers roles for IFT-A in retrograde train assembly and motility"

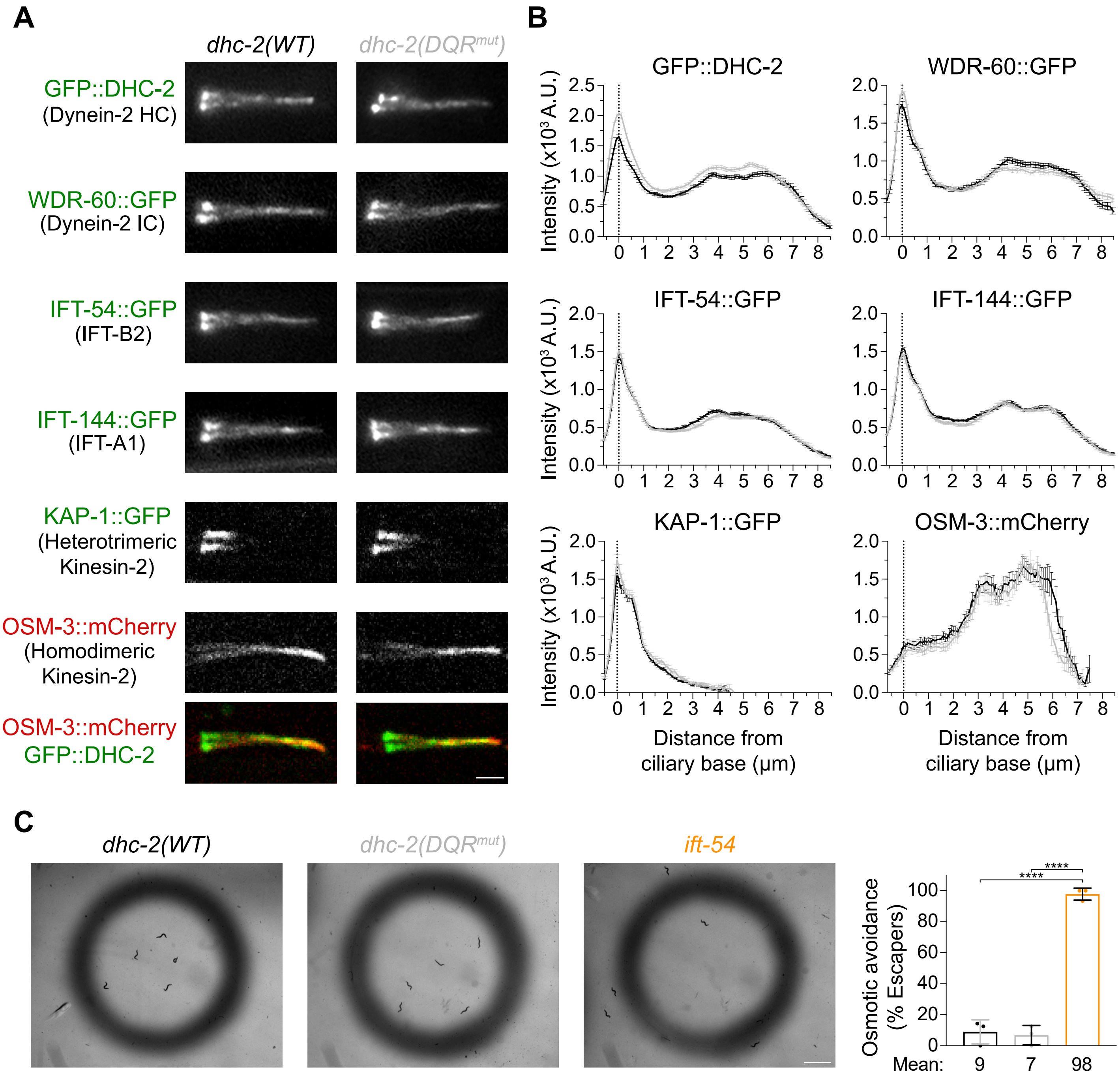


**Figure S1. IFT machinery distribution and ciliary function in dynein-2 DQR$^{mut}$ animals are identical to wild-type**

**(A)** Localization of multiple fluorescently-labeled IFT components in phasmid cilia from wild-type and dynein-2 DQR$^{mut}$ animals. **(B)** Intensity distribution profiles of the IFT components shown in *(A)* along cilia of wild-type and dynein-2 DQR$^{mut}$ genetic backgrounds (n≥66 cilia). **(C)** Osmotic avoidance assay to test dynein-2 DQR$^{mut}$ cilia function. *ift-54* mutant animals served as control given that their cilia are unable to detect the hypertonic glycerol barrier (n≥40 animals tracked over 8 separate assays). XY graphs show mean ±SEM; column graphs show mean ±SD. One-way ANOVA followed by the Dunnett multiple comparisons test was used to analyze the datasets in C. ****, P≤0.0001. Scale bars: 2 µm (A); 2 mm (C).

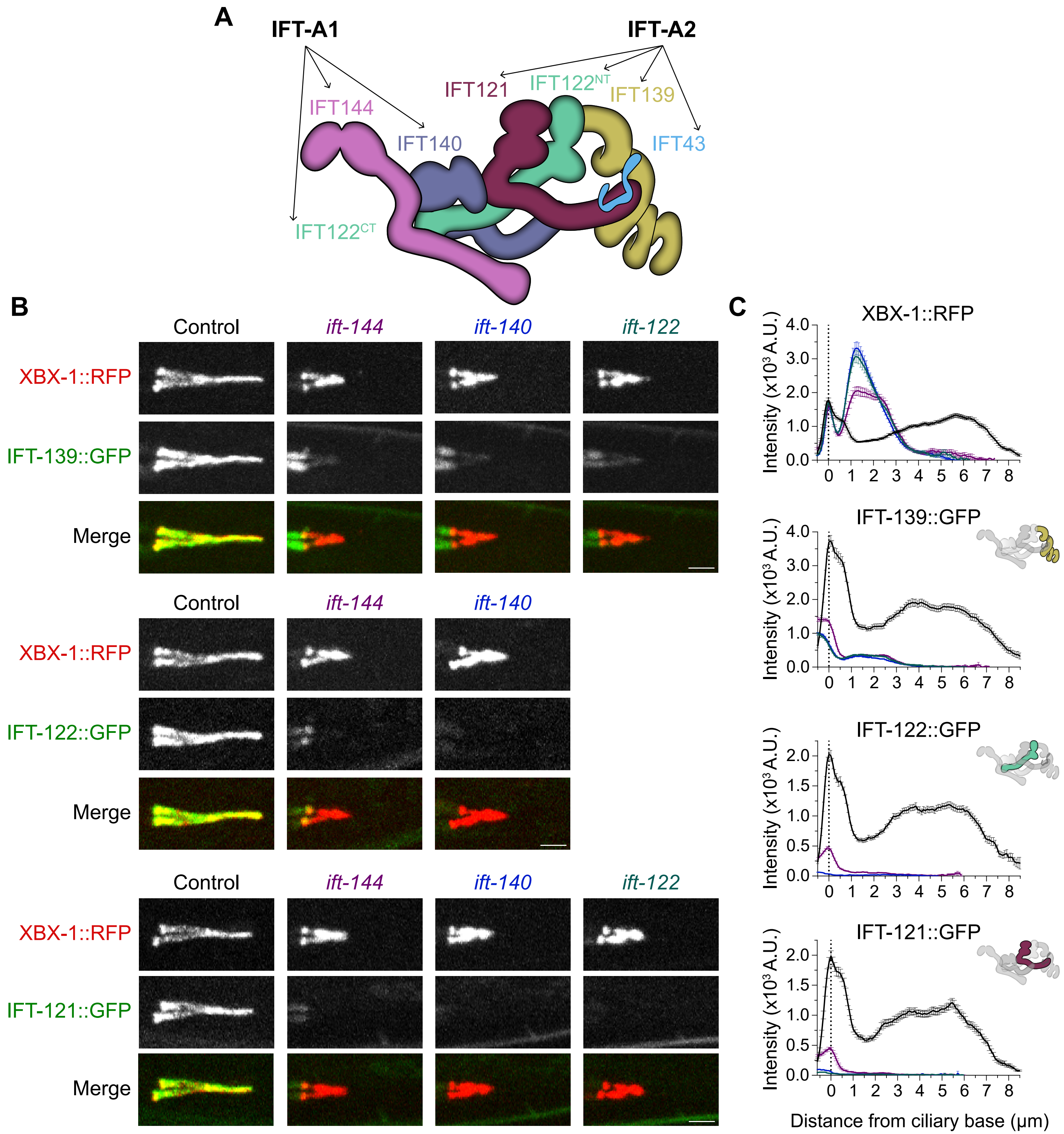


**Figure S2. IFT-A1 subunits are critical for IFT-A complex assembly and its incorporation into cilia**

**(A)** Illustration depicting the structure of the human IFT-A complex when docked into anterograde IFT trains, based in (Hesketh et al., 2022). This complex can be subdivided into the IFT-A1 and IFT-A2 modules (also known as IFT-A core and IFT-A peripheral modules, respectively). IFT-A1 is composed of IFT144, IFT140 and the C-terminal (CT) region of IFT122, while IFT-A2 encompasses IFT121, IFT139, IFT43, and the N-terminus (NT) of IFT122. **(B)** Localization of endogenously-labeled IFT-A2 subunits (IFT-139::GFP, IFT-122::GFP and IFT-121::GFP) in control and IFT-A1-deficient phasmid cilia, relative to the dynein-2 LIC (XBX-1::RFP). **(C)** Intensity distribution profiles of the IFT machinery components shown in *(B)* (n≥84 cilia). Similarly to GFP::DHC-2 and WDR-60::GFP, the LIC XBX-1::RFP strongly accumulated inside IFT-A-deficient cilia. The XBX-1::RFP distribution plot includes all the quantifications carried out in combination with the three GFP-tagged IFT-A2 subunits. Of note, while the IFT-122::GFP and IFT-121::GFP subunits were practically absent, a very small fraction of IFT-139::GFP was still incorporated into IFT-A1-deficient cilia. XY graphs are shown as mean ±SEM. Scale bars: 2 μm.

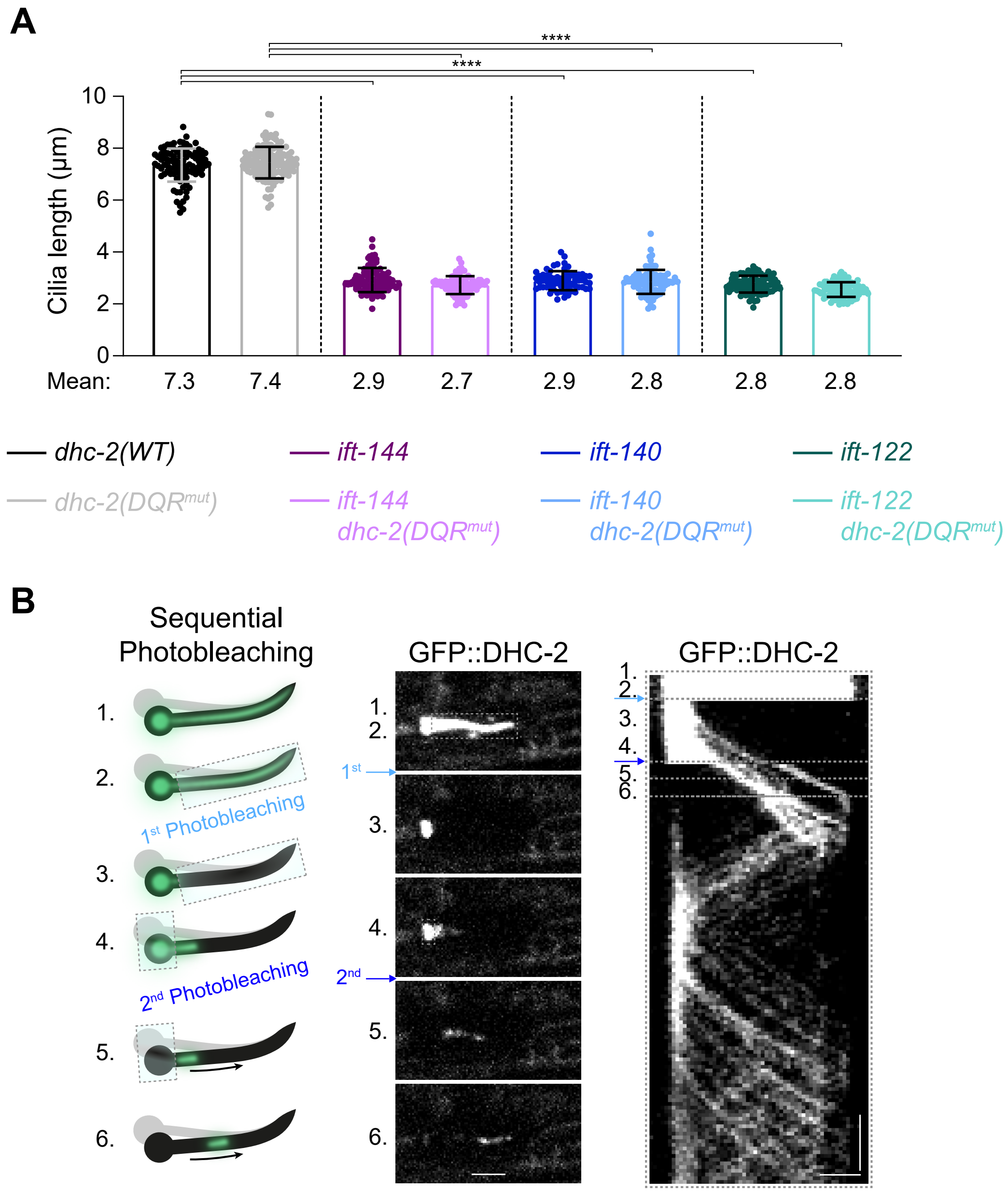


**Figure S3. Hot-wiring dynein-2 does not rescue axoneme extension in IFT-A1-deficient cilia**
**(A)** Average cilia length in IFT-A1 mutants, expressing the wild-type or DQR$^{mut}$ form of GFP::DHC-2. Data derived from size measurements carried out in the same cilia used to quantify the DHC-2 distribution profiles depicted in Figure 2B (n≥64 cilia). Data shown as mean ±SD. One-way ANOVA followed by the Dunn multiple comparisons test was used to analyze the datasets. ****, P≤0.0001. **(B)** Schematic detailing the sequential photobleaching technique (left) used in Figure 3F and 3G. Selected images (center) and the corresponding kymograph (right) from an example cilium illustrate the main stages of the protocol: 1) A phasmid cilium is brought into focus from the base to the ciliary tip; 2) The region comprising the ciliary axoneme is selected for the first photobleaching event, carefully avoiding the ciliary base; 3) Upon the first laser sweep, all IFT particles inside the cilium become photobleached; 4) A small fraction of IFT particles from the ciliary base is allowed to enter cilia, after which the ciliary base is selected for the second photobleaching event; 5, 6) The fraction of non-bleached IFT particles is then tracked along cilia, allowing for a more detailed view of IFT behavior in both control and mutant strains. Scale bars: vertical 5 s, horizontal 2 µm.

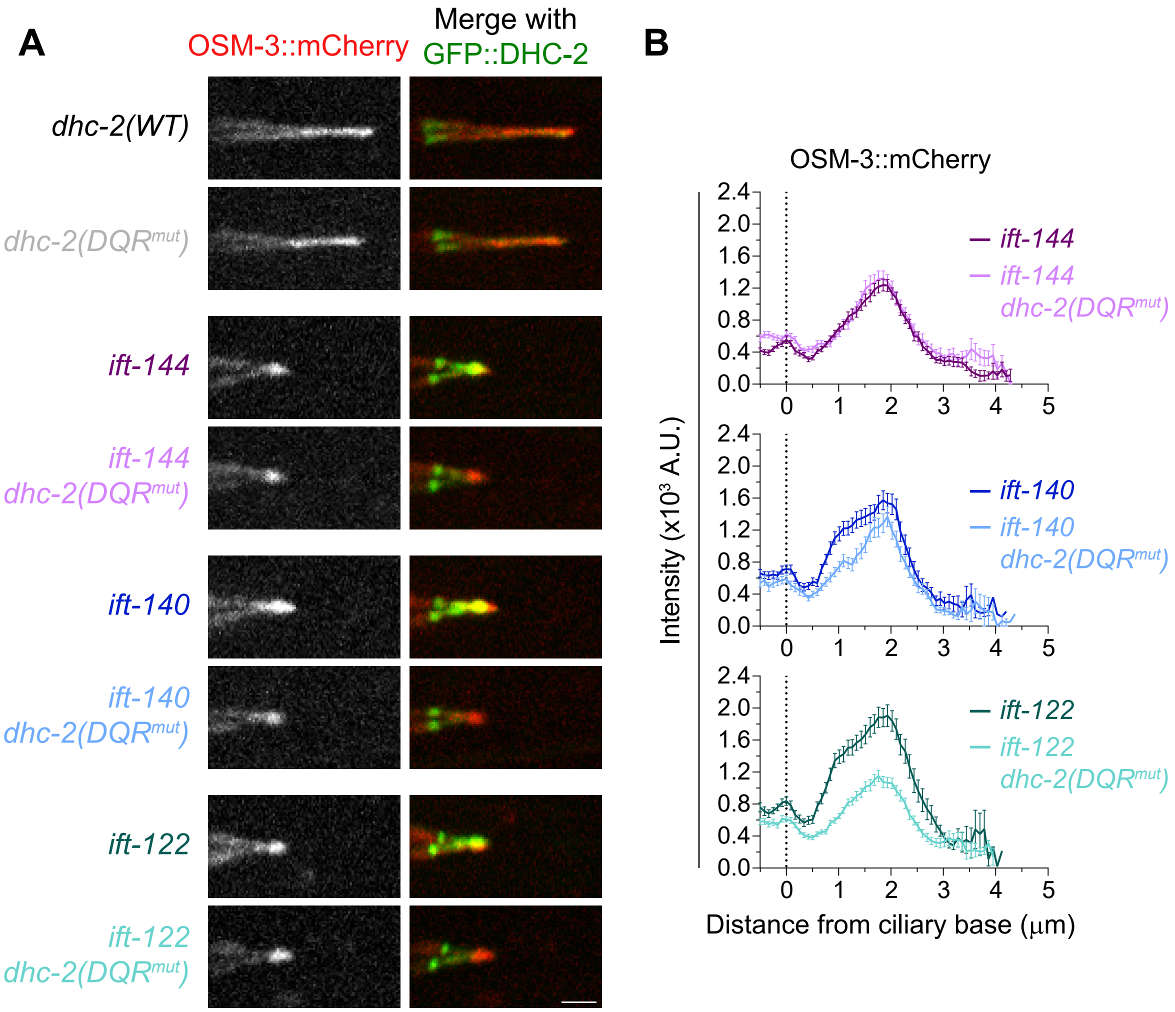


**Figure S4. Homodimeric kinesin-2 accumulates mostly at the tip of IFT-A1 mutant cilia**

**(A)** Phasmid cilia from control and IFT-A1 mutant strains, co-expressing OSM-3::mCherry and GFP::DHC-2 or GFP::DHC-2 $DQR^{mut}$. **(B)** Distribution profile of OSM-3::mCherry along cilia of the indicated genotypes (n≥ 56 cilia). XY graphs are shown as mean ±SEM. Scale bar: 2 μm.

**Table S1. Nomenclature of *C. elegans* proteins mentioned in the text and their corresponding orthologues in humans.**

| | Human proteins | *C. elegans* proteins |
|---|---|---|
| Dynein-2 Complex | DYNC2H1/DHC2 | CHE-3/**DHC-2** |
| | DYNC2I1/WDR60 | **WDR-60** |
| | DYNC2LI1/LIC3 | **XBX-1** |
| IFT-A Complex | WDR19/IFT144 | DYF-2/**IFT-144** |
| | IFT140 | CHE-11/**IFT-140** |
| | IFT122 | DAF-10/**IFT-122** |
| | TTC21B/IFT139 | **IFT-139** |
| | WDR35/IFT121 | IFTA-1/**IFT-121** |
| | IFT43 | **IFT-43** |
| IFT-B Complex | IFT74 | **IFT-74** |
| | TRAF3IP1/IFT54 | DYF-11/**IFT-54** |
| IFT Kinesin subunits | KIFAP3/KAP | **KAP-1** |
| | KIF17 | **OSM-3** |

Nomenclature used throughout the manuscript is highlighted in bold. Some alternative *C. elegans* gene nomenclature is used to facilitate its interpretation by readers working with other ciliated organisms. The alternative gene names were approved by WormBase curators prior to submission of this manuscript (changes will be made public in WormBase release WS289).
